# Season of transmission of Ross River/Barmah Forest Virus and *Mycobacterium ulcerans* closely align in southeastern Australia, supporting mosquitoes as the vector of Buruli ulcer

**DOI:** 10.1101/2023.08.07.552371

**Authors:** Andrew H. Buultjens, Ee Laine Tay, Aidan Yuen, N. Deborah Friedman, Timothy P. Stinear, Paul D.R. Johnson

## Abstract

Ross River Virus and Barmah Forest Virus infections (alphaviruses) have short incubation periods and are transmitted to humans by mosquitoes. *Mycobacterium ulcerans* infection (Buruli ulcer) has a much longer incubation period and its mode of transmission is contested. We studied the relationship between month of notification of alphavirus infections and Buruli ulcer in the temperate Australian state of Victoria over the six-year period, 2017-2022. Using *cross-correlation*, a signal processing technique, we found that a five-month temporal shift in month of Buruli ulcer notification provided optimal alignment with month of alphavirus notification. This closely matches the previously determined 5-month Buruli ulcer incubation period. Inferred transmission of both conditions showed coordinated maxima in summer and autumn and coordinated minima in winter and spring. The close alignment in season of transmission of alphavirus infection and Buruli ulcer in Victoria supports mosquitoes as the primary local vector of *M. ulcerans*.

## Introduction

Buruli ulcer (Buruli) is a geographically restricted, environmentally acquired infection caused by *Mycobacterium ulcerans* (1). Listed by WHO as a Neglected Tropical Disease, Buruli occurs in 33 countries but characteristically only in specific locations. Buruli can cause extensive tissue destruction if not diagnosed and managed effectively but recent advances in treatment with antibiotics have improved the outlook for sufferers in Buruli-endemic zones (2-7). The most active of these zones currently are west and sub-Saharan Africa (8), tropical Northern Australia (9) and coastal and urban zones of temperate southeastern Australia, particularly the state of Victoria (10).

In Victoria, an unprecedented outbreak of Buruli (figure 1) is ongoing with the disease now being transmitted in the inner suburbs of the state’s two largest cities of Melbourne and Geelong (11). The epidemiology of Buruli in Victoria has been shown to be zoonotic, centered on native possums with humans as accidental spillover hosts (12-14). The first evidence that mosquitoes may transmit *M. ulcerans* from possums to humans was published in this journal in 2007 (15, 16). This model of Buruli epidemiology is unusual and may be unique. Differences from the more familiar epidemiology in sub-Saharan Africa include Victoria’s coastal and urban endemic areas compared with inland rural riverine locations, our temperate climate compared with a tropical climate, a confirmed role for a small animal reservoir and a sustained rise in incidence compared with stabilization or decline in African endemic areas (17).

**Figure 1.**
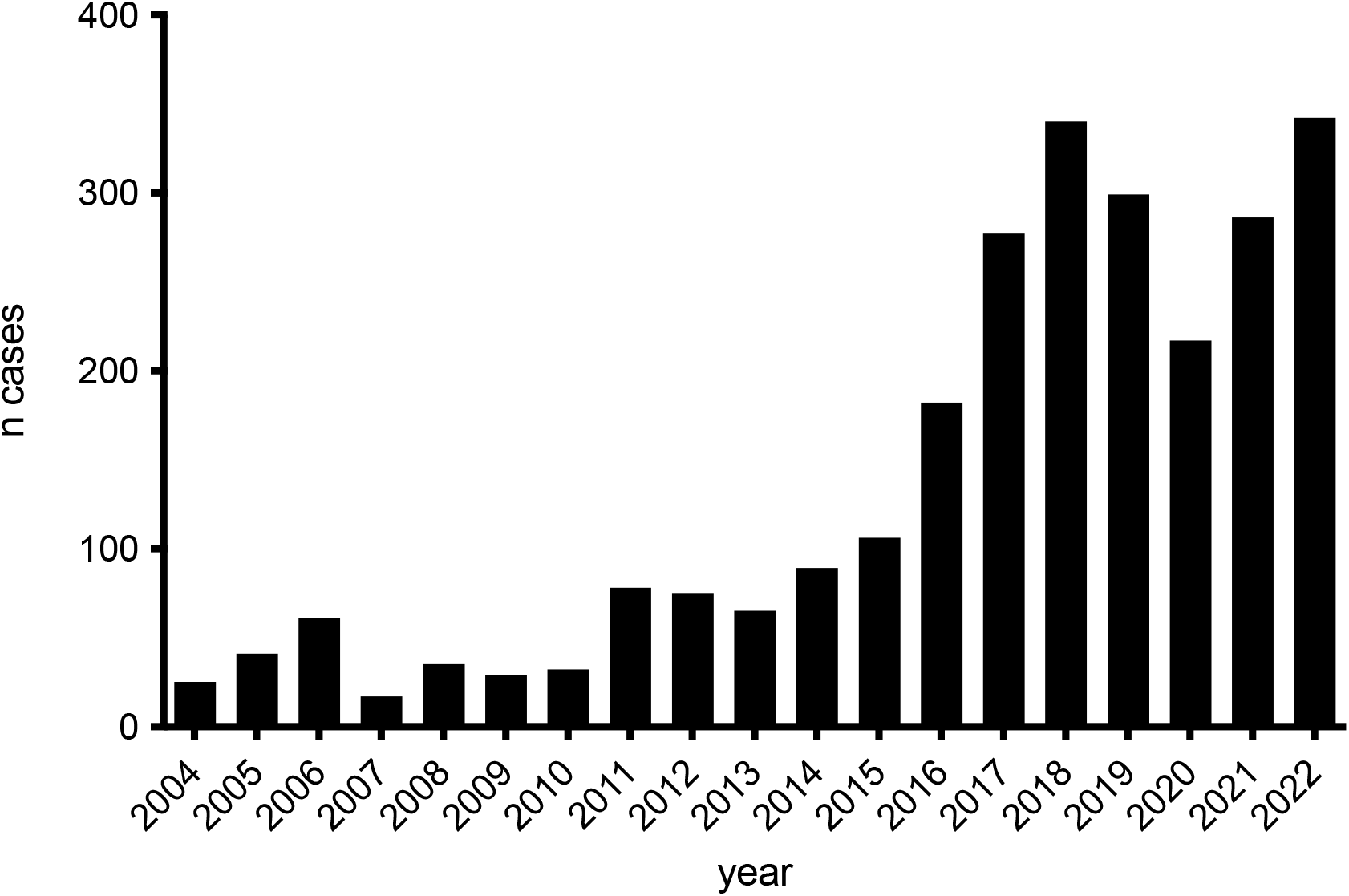
Victoria – n notified cases of Buruli by year.

Public health messaging in Victoria so far has been limited to health warnings to facilitate early diagnosis of Buruli and case mapping to inform local populations and primary care providers of the changing risk (18). To date there have not been coordinated programs aimed at Buruli prevention as the mode of transmission is contested.

Two reports published in this Journal in April 2009 (19) and December 2021 (20) considered the correlation between annual notification of alphavirus infections (Ross River virus and Barmah Forest virus) and Buruli in Victoria. These studies investigated the hypothesis that if the usual mode of Buruli transmission in Victoria is via mosquitoes, correlation with other conditions that are known to be mosquito-vectored would be expected. The first study identified a partial correlation between annual notifications of vector-borne diseases (alphavirus infections) and BuruIi in the calendar years 2002-2008 (18). However, in a subsequent report, Linke et al reported no ongoing statistical association since 2008 and concluded that factors other than mosquitoes were likely behind the change in Buruli incidence (20).

As the annual Buruli incidence in Victoria is now twenty times higher than it was when evidence for mosquito transmission was first published in 2007 (figure 1), we have revisited the correlation controversy using monthly rather than annual notification data and a novel statistical approach (cross-correlation). We hypothesized that a new analysis during a period of much higher Buruli incidence and a non-linear statistical approach could help resolve the transmission controversy with the hope that consensus can be reached.

## Methods

In collaboration with the Victorian Department of Health we accessed confirmed and probable notifications by month and year for Buruli and the two alphavirus infections – Ross River virus and Barmah Forest virus infections combined, over a 6-year period from 2017 to 2022.

Cases were defined according to national surveillance definitions for the two alphavirus infections (21, 22). Buruli was made notifiable in the state of Victoria from January 2004 due to local concerns about rising incidence. The surveillance definition for Buruli in Victoria has been recently published (23).

Month of notification for both Buruli and alphavirus infection was assumed to be the same as month of diagnosis.

Incubation periods for Buruli in Victoria have been calculated on two previous occasions by interviewing cases with only short exposures to known endemic areas but who lived and presented to doctors outside these areas. The median incubation period in Victoria was calculated to be between 4.5 (24) and 5 months (25). To calculate month of inferred transmission from month of notification for Buruli, the incubation period plus time to diagnosis/notification was assumed to be 5 months.

Incubation period for the two alphavirus infections is similar and has been reported as 3 days to 3 weeks, usually 1-2 weeks (26, 27). To calculate month of inferred transmission from month of notification, the incubation period plus time to diagnosis/notification for alphavirus infection was assumed to be 1 month.

Victoria has a temperate southern hemisphere climate with distinct seasons. Summer is defined as the months of December-February, autumn (fall) as March-May, winter as July-August and spring as September-November.

In addition to graphical representation, we examined the data with a signal processing technique called *Cross-correlation* to measure the relationship between the two epidemic curves (signals), as they are shifted in time. This avoids any bias or assumptions made in adjusting the data to obtain inferred transmission as it is based solely on the notification series itself. It also avoids assumptions about endemic and non-endemic area exposure which was the basis for estimations of the incubation period in previous work (24, 25).

To apply the method, incubation period adjusted alphavirus series (month of inferred transmission) and the raw Buruli ulcer notifications were used, so that the adjusted alphavirus signal may act as a temporal benchmark against which to align the Buruli ulcer notification series.

In the alphavirus time series, the first 3 months of 2017 were identified as outliers using z-scores (a statistical measure quantifying how many standard deviations a data point is from the mean), these timepoints were excluded from both datasets. The *correlate* function in the *numpy* python library (28) was used to identify the time shift factor in months that optimized the correlation between the two signals.

Data were displayed and analyzed using Graphpad Prism 10.0 (graphpad.com) or R (29).

## Ethics statement

Separate ethics approval is not required as data in this study were collected and used under the legislative authority of the Public and Wellbeing Act 2008 and only aggregated, de-identified data were used in this study. In addition, data by year and LGA are publicly available from the Department of Health (30). Data were summated by month and accessed with permission and assistance from senior epidemiologists at the Victorian department of Health.

## Results

Over the 6-year study period there were 3,839 alphavirus cases notified. Cases were strongly clustered by month and season and varied markedly from year to year (figure 2A). Notifications were much more likely in summer-autumn (3,503 notifications) compared with 336 in winter-spring, ratio 10.4:1. For Buruli there were 1,761 notifications and less variation across the 6 calendar years (figure 2). In marked contrast to alphavirus notifications, Buruli cases were 2.9 times more likely to be notified in winter-spring (1,307) than summer-autumn (454) (Table S1).

**Figure 2A.**
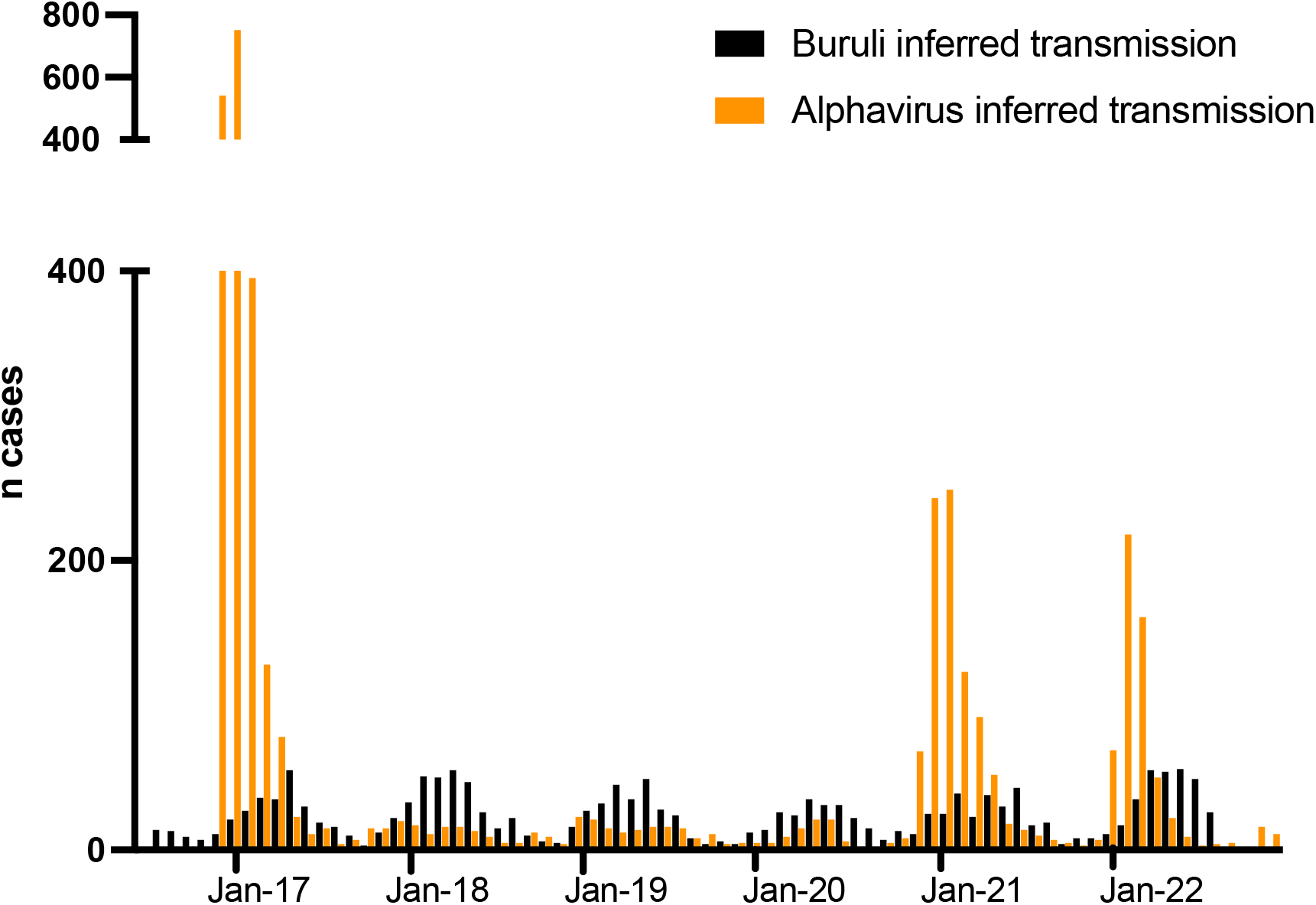
Month and year of notification for alphavirus infection and Buruli in Victoria. January of each year is the middle month of summer (x axis).

**Figure 2B.**
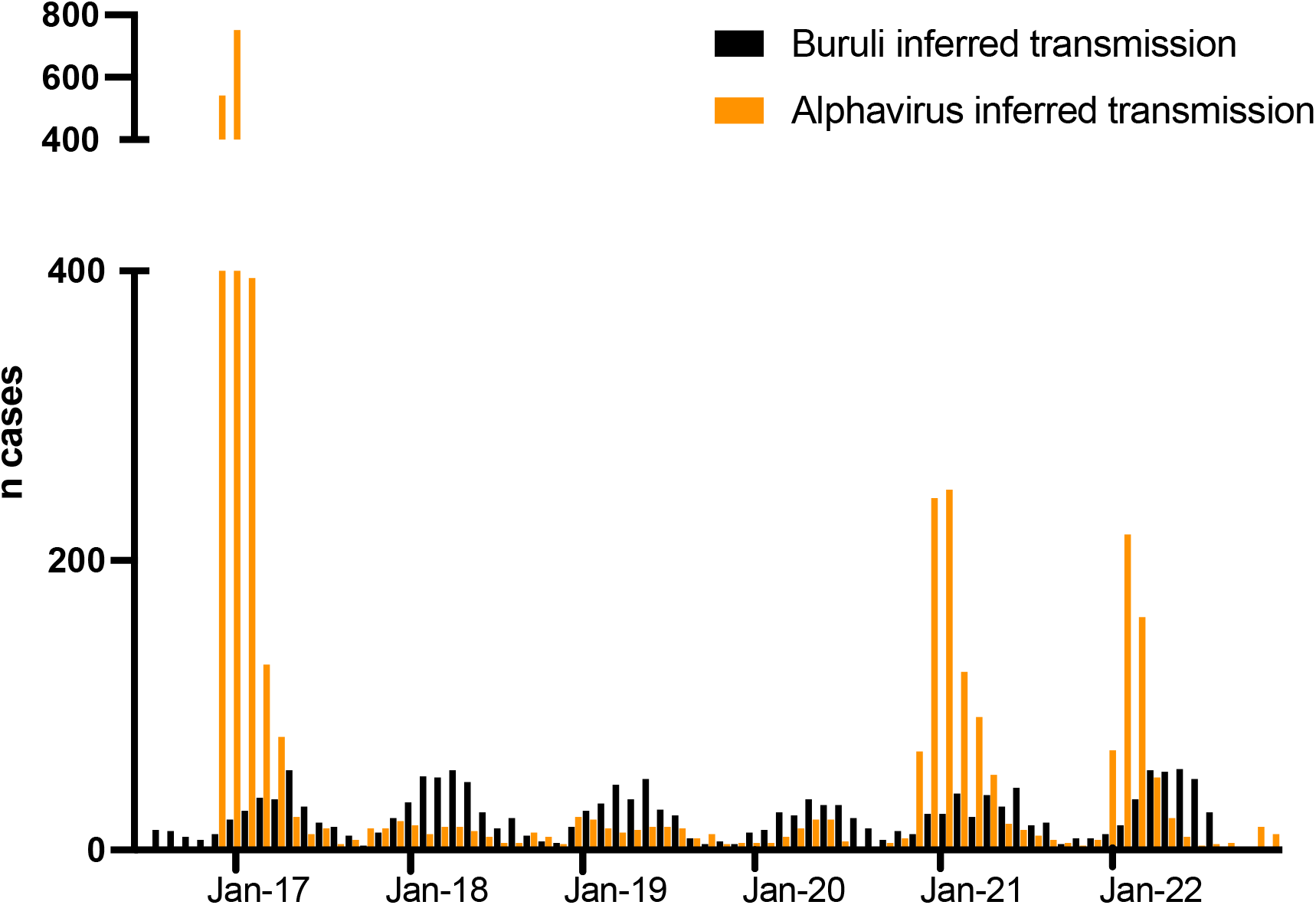
Month and year of notification manually adjusted to estimate inferred transmission. January of each year is the middle month of summer (x axis).

### Signal processing *cross correlation* analysis

To further explore the overlapping periodicity of the data, we employed a signal processing technique called *cross correlation* analysis. The two datasets were first inspected to identify and manage extreme outliers through a Z-score analysis (figure 3A). The initial three timepoints of the alphavirus time series were found to deviate more than three standard deviations from the mean, leading to their exclusion from both datasets (figure 3A).

**Figure 3.**
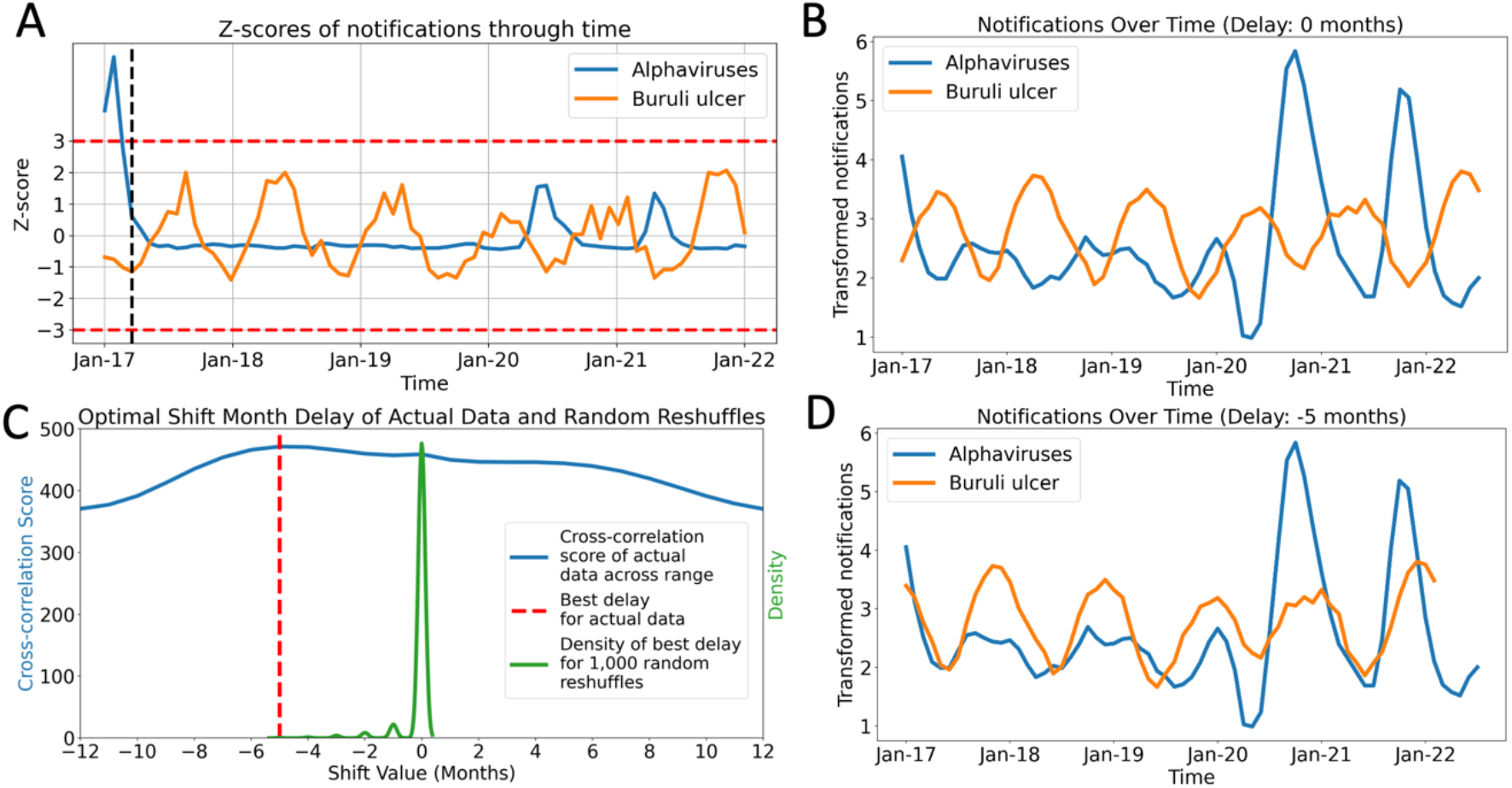
Analysis of Temporally Adjusted Buruli ulcer and alphavirus Notifications in Victoria (2017-2022). A) Z-score analysis of Buruli ulcer and Alphavirus Notifications. The horizontal red dashed lines represent the z-score boundaries of -3 and +3, which are equivalent to three standard deviations below and above the mean, respectively. The vertical black dashed line indicates the threshold used to exclude outlier data points falling beyond three standard deviations from the mean. B) Overlay of both time series without any temporal adjustment to the Buruli ulcer notification data. C) Cross-correlation analysis across a range of potential shift months (-12 to +12). The maximum correlation is observed with a shift of -5 months. D) Final alignment of the two datasets after the application of a -5-month shift to the Buruli ulcer notification data, optimizing their temporal alignment.

The censored data series then underwent logarithmic transformation and smoothing to amplify the cyclical signal within the data (figure 3B). The overlaid transformed series revealed an almost anti-phase relationship between the Buruli ulcer and alphavirus signals, indicating the presence of similar cyclic patterns that required temporal alignment for a better comparative analysis (figure 3B).

A cross-correlation analysis was then employed on the two series across a range of 48 shift months, which identified the peak cross-correlation score at -5 months (figure 3C). To establish the robustness of this optimal -5-month shift, a randomization analysis was conducted. This entailed performing the same cross-correlation analysis on 1,000 randomly reshuffled instances of the Buruli ulcer notification series. The resulting distribution from these iterations centered around zero, well separated from the optimal shift of -5 months (figure 3C).

When the Buruli ulcer notifications were then adjusted by this assumption-independent -5 months, a sinusoidal alignment with the alphavirus notifications was observed (figure 3D) that also very closely matched the previously established Buruli ulcer incubation period (24, 25).

## Discussion

There were 342 cases of Buruli notified in Victoria in 2022 compared with just 17 in 2007, a 20-fold increase since the year the first ever publications linking mosquitoes to Buruli transmission appeared in this journal (15, 16). Mosquito transmission was controversial and challenging to many when first published, although we had chosen to investigate this possibility after colleagues working in Africa had detected *M. ulcerans* in aquatic water insects in Buruli endemic areas (31, 32). In a research letter published in this Journal in April 2009 (19), we reasoned that if Buruli mosquito transmission was the usual mode of transmission in Victoria, year to year variation in Buruli notifications should be statistically associated with annual notifications of other vector-borne conditions (alphaviruses). We were able to demonstrate that over a 7-year period from 2002 to 2008 that Buruli and alphavirus annual notifications were correlated, albeit imperfectly. Subsequently, Linke et al re-investigated this association and found that the previously reported correlation was no longer present after 2008 using linear methods (20). They concluded that factors other than mosquitoes were likely behind the rising rates of Buruli in Victoria and that the lack of knowledge of the mechanism of disease transmission continues to hinder the implementation of successful public health interventions to control Buruli (20).

In this study, we revisited this controversy by examining monthly rather than annual notifications of Buruli and alphavirus infections and allowed for the effect of previously published incubation periods. Additionally, we identified the optimal temporal alignment of the notification datasets, without using prior knowledge of the previously estimated incubation periods for Buruli in Victoria (24, 25). This novel approach not only allowed us to identify potential correlations between the two conditions but also provides an independent estimate of the Buruli ulcer incubation period using only notification data without the need to interview cases or to make assumptions about which areas are endemic and which are not.

Our results show that inferred transmission of alphavirus infections and Buruli reach simultaneous maxima from December to May, and simultaneous minima from June to November every year over the 6-year study period. The accepted explanation in temperate Victoria for variation by season in alphavirus notifications is that warmer weather provides necessary climatic conditions for vectoring mosquitoes (33). Even though the animal reservoirs of alphaviruses are present throughout the year, transmission to humans falls almost to zero over the colder months.

A recently published detailed quantitative observational study showed that *M. ulcerans* in possum excreta and by inference in possum reservoirs is present in the environment throughout the year (13). Hence any hypothesis regarding Buruli transmission must account for this seasonal variation in transmission because environmental contamination remains constant despite the change in seasons. We believe that the simplest explanation is that transmission by mosquitoes is the usual way humans acquire Buruli in Victoria.

Interestingly, while we have shown a strong cyclic seasonal variation in inferred month of transmission for both Buruli and alphavirus infections, the magnitude of year-to-year variation is much greater for alphaviruses. Rainfall may partly explain the variation in alphaviruses as late 2016, and calendar years 2021 and 2022 were wetter and warmer in Victoria compared with historical averages, and 2018 and 2019 drier than historical averages (34-40). However, rainfall variation did not as strongly affect Buruli incidence. While we cannot currently explain this difference, we propose two hypotheses for future investigation. The first is that we know that for alphaviruses there is true biological vectoring by mosquitoes which brings with it exponential expansion in certain years. In contrast, while Buruli transmission is also strongly seasonally influenced, annual variation was less apparent possibly suggesting mechanical vectoring from a stable or only slowly changing pool of infected possums. Secondly, Buruli transmission in Victoria occurs mainly in areas with suburban style housing that is connected to potable water and where gardens are tended and protected from drought. Hence the impact of dry years on local mosquito and possum habitat may be reduced in Buruli endemic areas.

The limitation of the current study is that while we have shown a strong aligned seasonal effect of transmission risk for both alphavirus infection and Buruli in Victoria, we acknowledge that we have demonstrated correlation, not causation. However, there is a wealth of other published research supporting a central role for mosquito transmission of Buruli in Victoria. This includes evidence that human Buruli risk closely correlates with proportion of PCR-positive mosquitoes in 7 small towns in the Bellarine peninsula endemic area (41) and a new survey of mosquitoes demonstrating that 5.1/1000 of 65,000 mosquitoes tested PCR positive for *M. ulcerans* in the Mornington peninsula endemic area (42). A key new finding is confirmation that *M. ulcerans* cells detected by PCR in mosquitoes are the human outbreak strain (42). Also, along with our 2007 study where we reported detecting *M. ulcerans* by PCR at a rate of 4.2/1000 from 11,500 mosquitoes (15) we published a second paper also in 2007 describing a case control study investigating risk factors for Buruli acquisition on the Bellarine peninsula (16). In the final multivariate analysis, there were only two statistically significant factors remaining: reporting mosquito bites (odds ratio 2.56) and use of insect repellant (odds ratio 0.37). There was no significant association with bites from midges or march flies. Multiple other outdoor activities were examined including freshwater or saltwater swimming and fishing, surfing, sailing, bushwalking, lawn bowling, golf, bird watching, cycling, and gardening. None of these activities were interpedently associated with BU, suggesting that mosquito exposure specifically and not outdoor exposure generally increased the odds of Buruli acquisition.

We have attempted to investigate alternative modes of transmission including human skin temperature variation as an explanation for the non-random pattern of Buruli lesion distribution we observe (43). Our conclusion was that while there was a small effect, this was only a weak predictor of Buruli lesion distribution (44). We have also investigated the hypothesis that outdoor exposure in Buruli endemic areas could lead to transient skin contamination with *M. ulcerans* which could then be followed by a range of chance inoculating events including but not restricted to biting insects. We have not so far been able to find evidence to support this model (45).

In conclusion, the evidence that mosquitoes are the major vector of *Mycobacterium ulcerans* in temperate Victoria is very strong. Public health authorities responsible for Victorian Buruli endemic areas should now develop Buruli intervention programs that focus on mosquito bite prevention and mosquito control.

